# An improved molecular crowding sensor CRONOS for detection of crowding changes in membrane-less organelles under pathological conditions

**DOI:** 10.1101/2021.03.31.437991

**Authors:** Tamami Miyagi, Yoshiaki Yamanaka, Yuichiro Harada, Satoshi Narumi, Yuhei Hayamizu, Masahiko Kuroda, Kohsuke Kanekura

**Author notes:** equally contributed (Tamami Miyagi, Yoshiaki Yamanaka). Correspondence to: Kohsuke Kanekura, M.D., Ph.D. and Masahiko Kuroda, M.D., Ph.D., Kohsuke Kanekura, M.D., Ph.D, Assistant Professor, Department of Molecular Pathology, Tokyo Medical University, 6-1-1, Shinjuku, Shinjuku-ku, Tokyo, 160-8402, Japan., E mail, Masahiko Kuroda, M.D., Ph.D, Professor, Department of Molecular Pathology, Tokyo Medical University, 6-1-1, Shinjuku, Shinjuku-ku, Tokyo, 160-8402, Japan., E mail.

## Abstract

Membrane-less organelles (MLOs) formed by liquid-liquid phase separation (LLPS) play pivotal roles in biological processes. During LLPS, proteins and nucleotides are extremely condensed, resulting in changes of their conformation and biological functions. Disturbed LLPS homeostasis in MLOs cause fatal diseases such as amyotrophic lateral sclerosis. Therefore, it is important to detect changes of the degree of crowding in MLOs. However, it has not been investigated well due to lack of an appropriate method. To address this, we developed a genetically-encoded molecular crowding sensor CRONOS that senses the degree of macromolecular crowding in MLOs using fluorescence resonance energy transfer (FRET) system. CRONOS is a very bright biosensor with wider dynamic range and detect changes in the macromolecular volume fraction better than the previously reported mCer-mCit sensor in solution. By fusing to scaffold protein of each MLO, we successfully delivered CRONOS to MLO of interest and detected previously undescribed difference of the degree of crowding in each MLO. If not tagged, CRONOS localized to interstitial space of MLOs, giving us the crowding information of inspace. CRONOS also detected changes of degree of macromolecular crowding in nucleolus induced by environmental stress or inhibition of transcription. These findings suggest that CRONOS can be a useful tool for determination of molecular crowding and detection of pathological changes in MLOs in live cells.

## Introduction

Eukaryotic cells contain extremely high concentrations of proteins at 50-400 mg/ml, and are filled with various nucleic acids and macromolecules in addition to proteins^12^. Such high concentration of proteins in cells are crucial for the efficient biochemical reactions^3^. In order for these molecules to coordinately maintain vital activities while avoiding disorderly mixing, it is efficient for biomolecules to form local compartments. The examples of classical compartments are intracellular membranous organelles such as mitochondria and endoplasmic reticulum partitioned by lipid membranes. The membranous organelles are suitable for sustaining constitutively specific biochemical reactions in an isolated environment. On the other hand, it has been reported that molecules that are not captured in the membrane organelles transiently undergo liquidliquid phase separation (LLPS) and form membrane-less organelles (MLOs), which is preferable for transient biochemical reactions in response to various stimuli^4^. The biological roles of MLOs are highly diverse: nucleolus is the site for ribosome biogenesis and protein quality check under stress conditions^5,6^; stress granules control mRNA stability and protein translation under stress conditions^7^; nuclear speckles are the site for storage and modification of splicing factors^8^. MLOs consist of scaffold proteins specific for each MLO and nucleic acids, and these proteins and nucleic acids undergo LLPS to form each MLO^9,10^. During the phase-separation process, the scaffolding proteins and nucleic acids are highly condensed in the droplet through the biophysical forces such as electrostatic force, cation-pi interaction and hydrophobic interaction derived from its component proteins and nucleic acids^11,12^. It is estimated that the concentration of specific proteins which undergoes phase-separation can be up to more than hundreds-fold in the droplets compared with the concentration in the supernatant fraction^13^. Such a high density of specific species of proteins can alter the fashion of protein-protein interaction, reaction kinetics, protein conformation and overall functions. In addition, disturbance of LLPS by pathological changes of the composing proteins result in impairment of LLPS homeostasis and malfunction of MLO, causing fatal diseases including amyotrophic lateral sclerosis (ALS)^14,15^.

Although MLO play pivotal roles in a wide range of biological phenomena, there are many uncertain points to be clarified. For example, it remains unknown whether the degree of crowding varies in each type of MLO. Furthermore, it is unclear how the crowding of MLO is affected by biological processes, due to lack of appropriate tools to measure them. The first attempt to directly measure macromolecular crowding in cells was achieved by the FRET-based biosensor with mCerluean-mCitrine (mCer-mCit)^16^. The sensor successfully detected the crowding change in cells induced by osmotic shock. However, its relatively small dynamic range might not be enough to assess subtle physiological changes.

To investigate macromolecular crowding especially in MLOs in live cells, we developed a FRET-based biosensor CRONOS (<u>cro</u>wding sensor with m<u>N</u>e<u>o</u>nGreen and m<u>S</u>carlet-I) which has a wider dynamic range than the previously reported mCer-mCit sensor. CRONOS successfully detected the degree of macromolecular crowding of various MLO in live cells under normal and pathological conditions.

## Materials and Methods

### Crowding sensor design

The fluorescent protein-based FRET sensor was designed based on the previously reported molecular crowding sensor with a FRET pair of mCerulean-mCitrine (mCer-mCit)^16^. The sensor consists of mCer as a FRET donor and mCit as a FRET acceptor, sandwiching a linker consisting of α-helix forming sequences. We sought for a better FRET-pairs for the sensor and found that the mNeonGreen-mScarlet-I (mNG-mScaI) pair would work better^17^. CRONOS (<u>cro</u>wding sensor with m<u>N</u>e<u>o</u>nGreen and m<u>S</u>carlet-I) consists of mNG as a FRET donor and mScaI as a FRET acceptor, sandwiching the same linker which is used in mCer-mCit sensor. The cDNAs encoding mCer-mCit sensor or CRONOS were synthesized and subcloned into pET28a vector containing His6 tag (Genscript, NJ, USA).

### Purification of recombinant sensor proteins

E. coli BL21 strain was transformed with pET28a-mCer-mCit sensor or pET28a-CRONOS and plated on a LB-agar plate containing ampicillin, incubated for overnight. Single colony was picked up for each construct, and after culturing overnight at 37°C, expression of recombinant proteins was induced by incubation with 0.1 mM isopropyl β-D-1-thiogalactopyranoside (IPTG) for 3 hr at 30°C. The recombinant proteins were purified with His60 Ni Gravity Column Purification Kit (Clontech) following a manufacturer’s protocol. After elution by 300 mM imidazole, the recombinant proteins were dialyzed with phosphate-buffered saline (PBS) using Slide-A-Lyzer dialysis cassette (ThemoFisher) for 4 hr. The purified proteins were then frozen until use.

### *In vitro* experiments

Both mCit-mCer sensor and CRONOS were challenged with crowding macromolecules, polyethylene glycol (PEG) of various molecular weights, Ficoll PM400 and D-glucose dissolved in PBS at different concentration (5-40% weight/volume) as indicated. Each sensor protein was added to the solution at final 2-3 μg/ml otherwise mentioned. After mixing the solution and sensor, the FRET efficiency was assessed by RF-6000 fluorescence spectrophotometer (Shimazu, Kyoto, Japan). For mCer-mCit sensor, the donor mCer was excited at 433 nm and the emission spectra was measured. For CRONOS, the donor mNG was excited at 506 nm and the emission spectra was measured. To calculate the FRET ratio of mCer-mCit sensor, the fluorescence intensity of 527 nm was divided by the fluorescence intensity of 476 nm. To calculate the FRET ratio of CRONOS, the fluorescence intensity of 588 nm was divided by the fluorescence intensity of 522 nm.

### LLPS in solution

The LLPS model in solution was made by (PR)_12_ peptide and poly-rA RNA (Roche) in water or PBS^18^. (PR)_12_ peptide was chemically synthesized with purity higher than 85%, followed by removal of trifluoroacetic acid (Genscript). When 100 μM of (PR)_12_ peptide and 0.5 mg/ml of poly-rA RNA were mixed, they underwent LLPS in water or in PBS. After mixing the solutions, the sensors were added.

### Cellular experiments

For cytosolic expression in mammalian cells, cDNAs encoding mCer-mCit sensor or CRONOS were subcloned into pcDNA3.1(+) vector. For targeting the sensor to each MLO, we cloned cDNA of each MLO marker protein by PCR with HeLa cell cDNA, and the CRONOS was fused to N-terminus of human NPM1 (nucleolus) or N-terminus of human TIA1 (stress granule). HeLa cells were cultured in Dulbecco-modified Eagle’s Medium (DMEM) supplemented with 10% fetal bovine serum (FBS) and antibiotics, maintained in a standard cell culture condition. For imaging, HeLa cells, plated on a chambered coverglass (Matsunami) or a chambered slideglass (Matsunami), were transiently transfected with pcDNA3.1-CRONOS derivatives using Lipofectamine 2000 reagent (ThermoFisher). The cells were challenged with heat shock at 42 °C for 3 hr, or 2 μg/ml of Actinomycin D treatment for 3 hr. The cells were imaged with LSM710 laser confocal microscopy using 63× water immersion objective lens (Zeiss) approximately 24 hr after transfection. CRONOS was imaged by exciting the donor mNG at 488 nm and collecting two simultaneous images of the donor mNG (520-560 nm) and the acceptor mScaI (590LP) at same gain settings. Image ratio calculation was performed with Fiji software (www.imagej.net). Fluorescence loss in photobleaching (FLIP) analysis was performed by LSM710 with Zen software (Zeiss). For measurement of fluorescence emission spectra in live cells, HEK293 cells were transiently transfected with the sensor plasmids. Twenty-four hr after transfection, the cells were pelleted and resuspended in PBS or PBS with 400mM sorbitol at 3 million cells/ml and the spectra were measured by RF-6000 fluorescence spectrophotometer (Shimazu).

### Statistics

All data are represented as mean ± standard deviation. Statistical analysis was performed with SPSS software 26 (IBM).

## Results

There are several types of molecular crowding sensors using FRET system such as fluorescent protein-based sensors and nucleic acid-based sensors^16,19,20,21^. Because we intended to investigate the degree of crowding in MLOs, we needed to control delivery of the sensor to specific MLO of interest. For this end, protein-based sensor has an advantage to add organelle-targeting signal sequences. The previously reported mCer-mCit crowding sensor consists of N-terminal mCer (cyan fluorescent protein) and C-terminal mCit (yellow fluorescent protein) sandwiching two α-helical peptides with a hinge which works as a conformationally flexible linker (Figure. 1A), and the sensor could detect cellular response to osmotic stress induced by sorbitol in live cells^16^. The physiological changes of macromolecular crowding in MLOs might be more subtle than the changes induced by chemical molecular crowders and thus we sought for a better FRET pair which confers a wider dynamic range to the sensor. From the literature search, we chose the FRET pair with best performance^17^ consisting of mNG (green fluorescent protein: ex. 506 nm/em. 517 nm)^22^ as a donor protein and mScaI (red fluorescent protein: ex. 569 nm/em. 593 nm)^23^ as an acceptor protein, and named the sensor CRONOS (<u>cro</u>wding sensor with m<u>N</u>e<u>o</u>nGreen and m<u>S</u>carlet-I) (Figure. 1B)^17^. Both mNG and mScaI work as a monomer with minimal self-association^22,23^. When we challenged these sensors to a macromolecular crowding reagent, 20%-PEG8000, both CRONOS and the mCer-mCit sensor successfully detected the crowding (Figure. 1C and 1D). To deny the possibility that the change was caused by self-associated inter-molecular FRET between multiple sensor molecules due to high concentration, we titrated the concentration of CRONOS from 10 nM to 200 nM, and we confirmed the FRET ratio was not changed by concentration of the sensor (Figure. 1E and 1F). The strong signal of CRONOS, derived from very bright natures of mNG and mScaI, also enables various detection methods (Figure. 1G).

**Figure 1.**
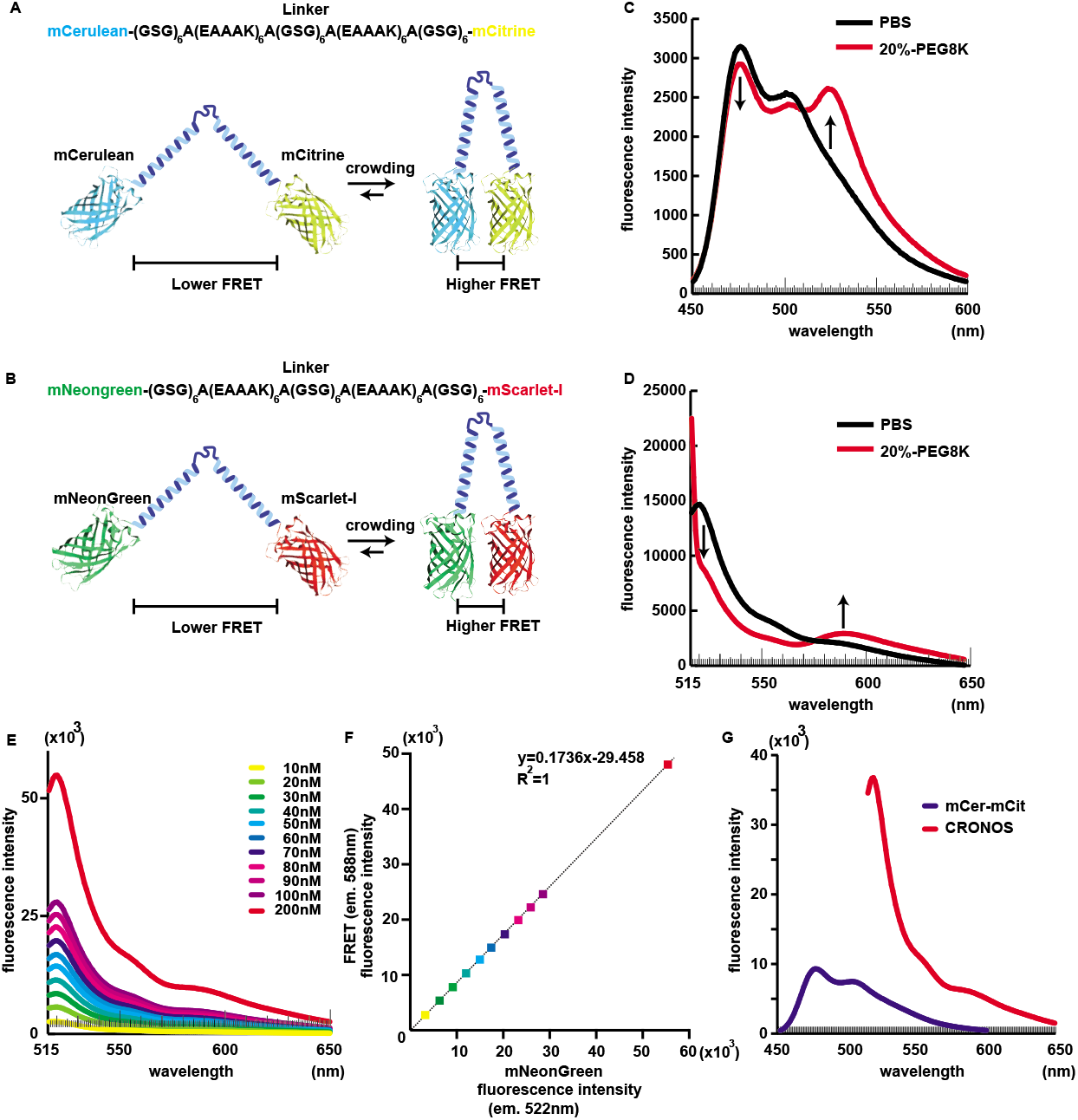
Optimization of FRET pair produces a bright protein-based crowding sensor. (A) (B) The scheme shows design and concept of crowding sensor based on mCer-mCit FRET system (A) and CRONOS FRET system (B). The FRET donor and acceptor are connected via a conformationally flexible linker composed of two α-helical peptides with a hinge. Upon molecular crowding, the sensor confirmation gets more condensed, resulting in higher FRET efficiency. (C)(D) Fluorescence emission spectra of mCer-mCit sensor (C) or CRONOS (D) in PBS or in 20% PEG8000-PBS revealed that molecular crowding by PEG increased FRET efficiency of both sensors. The template structure of fluorescent protein was obtained from GFP (4EN1) on Protein Data Bank (PDB). (E) (F) The effect of protein concentration on the fluorescence emission spectra (E) and the FRET ratio (F). (G) Comparison of fluorescence intensity of mCer-mCit sensor and CRONOS sensor. Each sensor at 100 nM was measured.

Next, we performed detailed comparison of the two sensors in solutions. When we excited the mCer-mCit sensor at 433 nm, we observed a basal FRET ratio (mCit/mCer: 527 nm fluorescence / 476 nm fluorescence) of 0.51 (Figure. 2A and 2B). Titration of the mCer-mCit sensor with PEG300 or PEG8000 resulted in increase of FRET ratio (1.14-fold increase for 40% (w/v) solution of PEG300, 2.86-fold increase in 40% solution of PEG8000, respectively) but D-glucose did not affect FRET ratio (Figure. 2A, 2B and Figure. S1A, S1B). Next, we examined CRONOS. To avoid leak of excitation light, we used fluorescence intensity at 522 nm as mNG signal hereafter. When we excited the donor mNG at 506 nm, we observed a basal FRET ratio (mScaI/mNG: 588 nm fluorescence / 522 nm fluorescence) of 0.17 (Figure. 2C and 2D). Titration of CRONOS with PEG300 or PEG8000 resulted in increase of FRET ratio (1.57-fold increase in 40% solution of PEG300, 9.0-fold increase in 40% solution of PEG8000, respectively) but D-glucose did not increase FRET ratio either (Figure. 2C, 2D and Figure. S1C, S1D), showing that CRONOS have a wider dynamic range to measure macromolecular concentration.

**Figure 2.**
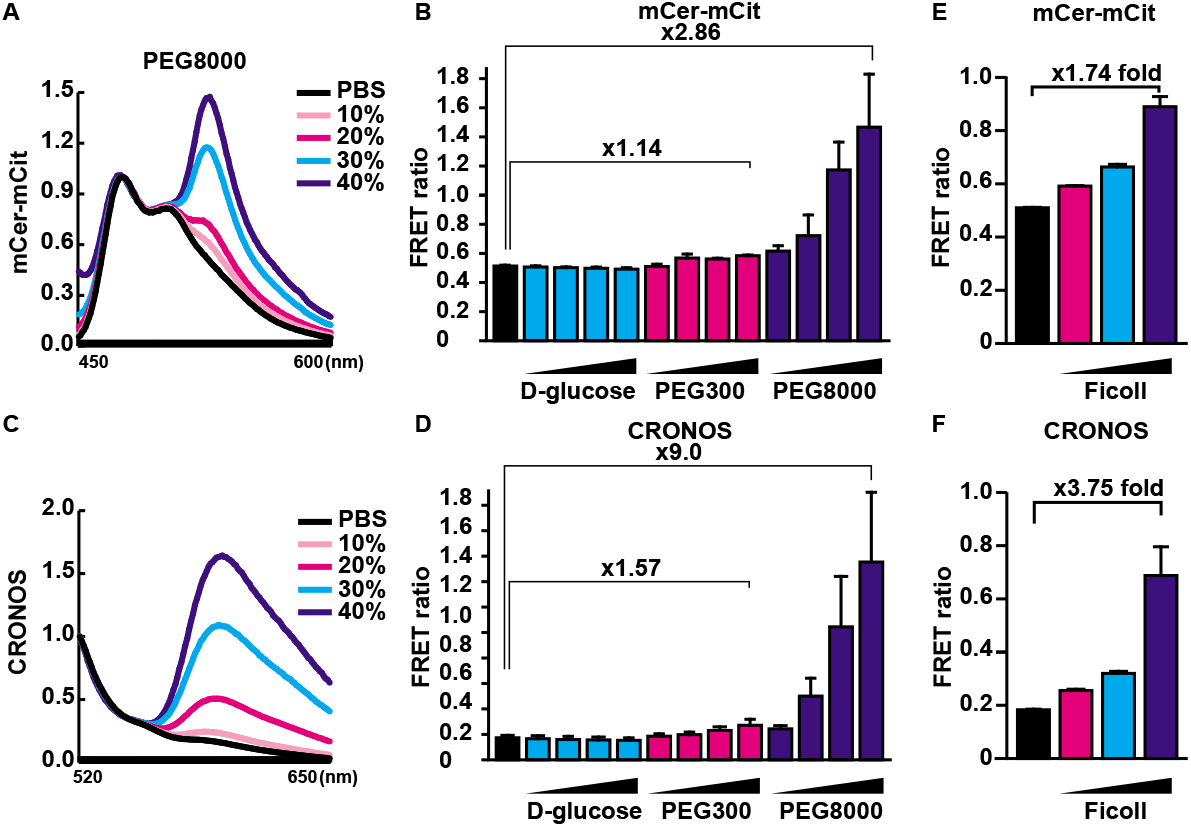
CRONOS detects molecular crowding change in solutions. (A) Fluorescence emission spectra of the mCer-mCit sensor upon titration with 10%-, 20%-, 30%- or 40% of PEG8000 were shown. The fluorescence emission spectra were normalized to the donor fluorescence. (B) The FRET ratio of the mCer-mCit sensor exposed to 10%-, 20%-, 30%- or 40% of D-glucose, PEG300 or PEG8000 were shown. (C) Fluorescence emission spectra of CRONOS upon titration with 10%-, 20%-, 30%- or 40% of PEG8000. (D) The FRET ratio showed better performance of CRONOS in response to PEG300 and PEG8000. (E) The mCer-mCit FRET ratio upon treatment with PBS containing 5%-. 10%- or 20% of Ficoll PM400. (F) The FRET ratio showed better performance of CRONOS in response to Ficoll PM400. The error bars indicate ±S.D. obtained from three independent measurements.

We next examined the effect of volume exclusion effect of different size of PEG. When the molecular weight of PEG increased, the volume exclusion effect also gets greater, suggesting that the bending of sensor efficiently occurs. When we mixed the two sensors with 10% of PEG with different molecular weight, the change of FRET ratio was bigger in CRONOS than in mCer-mCit (Figure S1E-S1H), also substantiating the idea that CRONOS have a wider dynamic range. We also measured the change of FRET ratio by Ficoll, and CRONOS showed better performance with Ficoll (Figure 2E, 2F and Figure S1I, S1J).

Macromolecular condensation is the biophysical principle for formation of MLOs in cells, and condensation of disease-causing proteins are widely studied to clarify pathological changes of phase-separated proteins^13,24^. To test if CRONOS can detect change in crowding in phase separate droplets, we tested these biosensors in LLPS in solutions. For this end, we adopted a model of biomolecular condensates consisting of (Pro-Arg: PR)_12_ dipeptide encoded by C9orf72 gene, a leading cause of familial amyotrophic lateral sclerosis (ALS), and poly-A RNA which are reportedly undergo LLPS when mixed. Upon mixing of (PR)_12_ and poly-A RNA in solution, highly cationic dipeptides and highly anionic RNA interact each other, undergo LLPS and form droplets. To assess the molecular crowding in the droplets, the sensor must be uptaken by the LLPS droplets. The surface of poly(PR) and RNA droplet is negatively charged due to the z potential ^25^, and this negative charge is supposed to attract the sensor molecules to the droplets. First, we added recombinant mCer-mCit sensor to water, poly-rA, (PR)_12_ peptide and the mixed LLPS droplets dissolved in water. The FRET ratio of mCer-mCit was increased by (PR)_12_, but not by LLPS droplets (Figure S2A). We observed the solution with confocal microscopy and found that mCer-mCit sensor formed aggregates with (PR)_12_ and weakly deposited on the surface of phase-separated droplets (Figure S2B). We next tested recombinant CRONOS. The FRET ratio of CRONOS was decreased by poly-rA and (PR)_12_ whereas increased in the LLPS droplets. When we tried these sensors in PBS-based solution, the changes of FRET ratio of both mCer-mCit sensor and CRONOS were blunted (Figure. 3A and Figure S2A). When observed the solution with confocal microscopy, the uptake of mCer-mCit sensor and CRONOS to the droplets was strongly inhibited by PBS, indicating the limited usefulness of these sensors in LLPS droplets in solutions (Figure. 3B and Figure S2B). These results indicated that when attempting to measure the crowding in LLPS droplet, tagging the sensor to phase-separating molecules will be better than spontaneous uptake of sensor by electrostatic force, and when using the sensor in solutions, we need to pay attention to salt concentration and distribution of sensor molecules.

**Figure 3.**
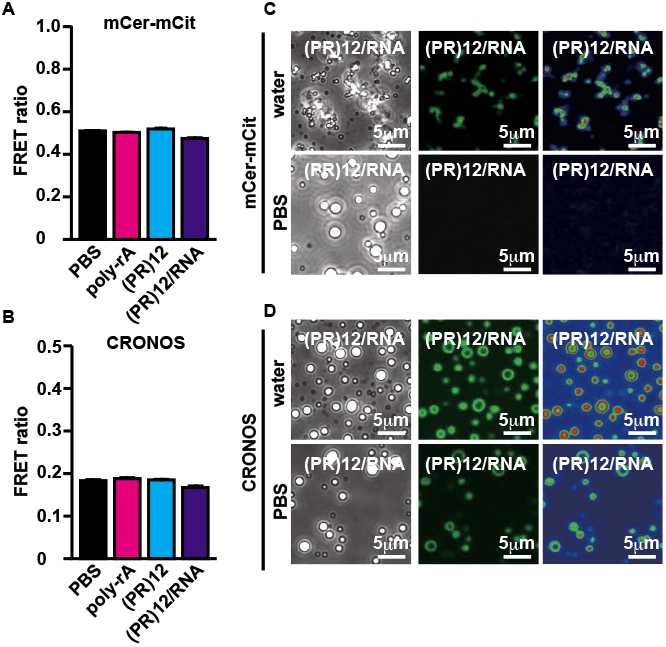
Uptake of crowding sensors into phase-separated droplets is insufficient in water, and further inhibited by existence of salt. (A) Neither mCer-mCit sensor or CRONOS detected molecular crowding in LLPS droplets in PBS. (B) Existence of salt attenuated uptake of the sensor into the phase-separated droplets. Right panels were pseudo-colored images of GFP channel. The error bars indicate ±S.D. obtained from three independent measurements.

Next, we tested whether CRONOS can detect the macromolecular crowding in live cells, especially in MLOs. For this end, we made three CRONOS constructs, CRONOS (without targeting sequence), CRONOS-NPM1 (nucleolus-targeted) and CRONOS-TIA1 (stress granule-targeted). In previous studies, both of the protein-based crowding sensor and nucleic acid-based sensor indicate that cytosolic macromolecular density is higher than that of nucleus ^19,20^. When we overexpressed CRONOS without targeting sequence in HeLa cells, it diffusely located to cytosol and nucleus, and a small portion of CRONOS distributed to nucleolus (Figure. 4A). The FRET ratio of mScaI/mNG showed that nuclear crowding was lower than that of cytosol as reported, and nucleolar FRET ratio was the lowest (Figure. 4A). On the other hand, CRONOS-NPM1 mainly distributed to nucleolus and a small portion of the sensor located to nucleoplasm, and nucleolar FRET ratio was higher than that of nucleoplasm (Figure.4A) which was opposite from the result from CRONOS without tag. Overexpression of TIA1 is reported to form stress granules, and indeed CRONOS-TIA1 formed stress granule like dots in cytosol with the highest FRET ratio among organelles measured (Figure.4A). These data showed that molecular crowding varies among MLOs.

**Figure 4.**
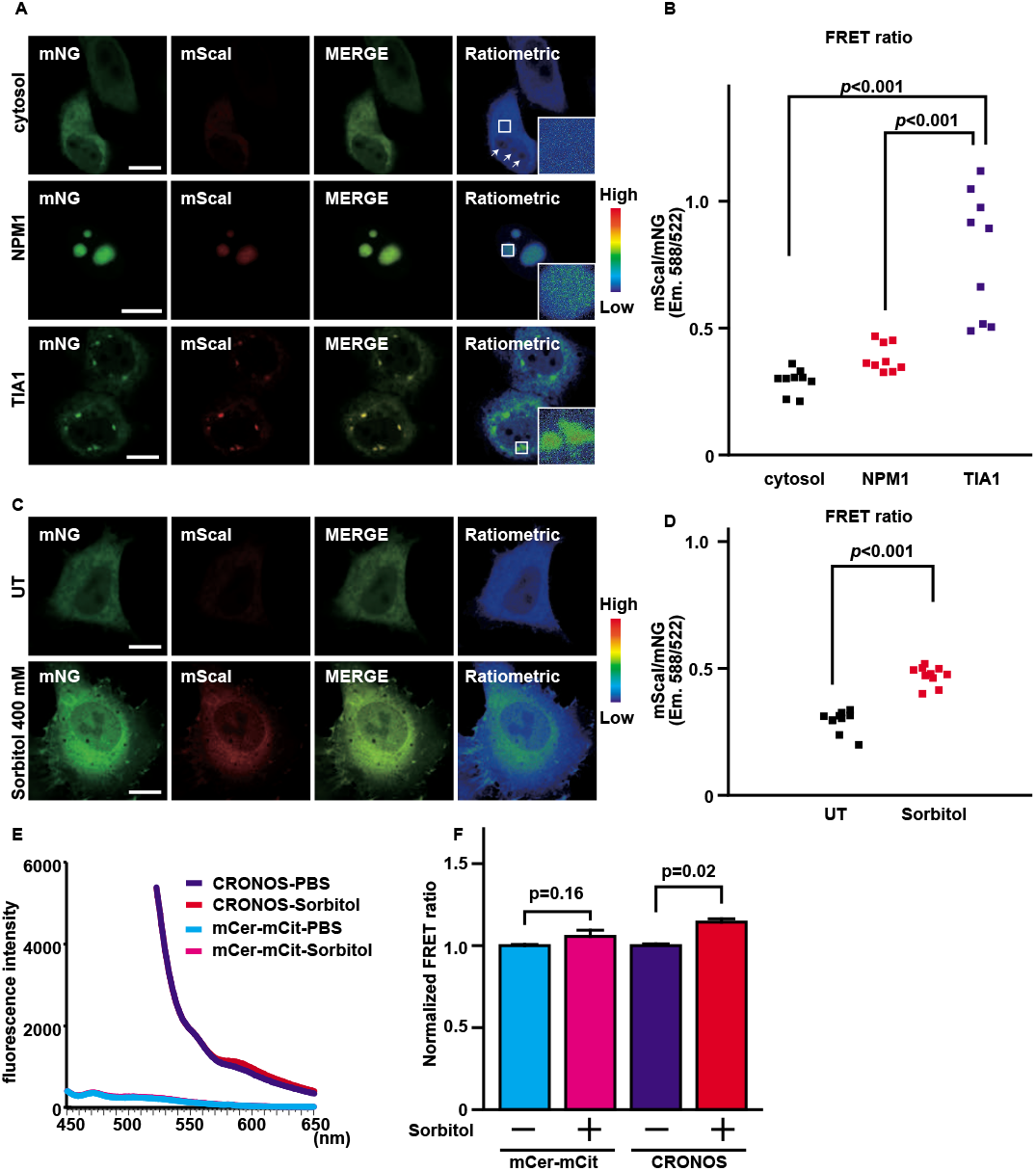
CRONOS sensor detects the changes of crowding in different organelles. (A) (B) HeLa cells overexpressing CRONOS without tag (cytosol), CRONOS-NPM1 (nucleolus) or CRONOS-TIA1 (stress granule) were imaged by a confocal microscopy. The cytosol, nucleolus and stress granules were imaged with same setting, enabling the comparison of their crowding. Whiter arrows showed nucleoli. The white squared areas were zoomed. (C)(D) HeLa cells overexpressing CRONOS treated with or without 400 mM sorbitol. (E)(F) Emission spectra and FRET ratio of mCer-mCit sensor and CRONOS in live HEK293 cells. The scale bar shows 10 μm.

To test whether CRONOS can detect dynamic change of molecular crowding, we challenged the HeLa cells with 400 mM sorbitol for 10 min to induce osmotic stress, causing cytosolic condensation. CRONOS successfully detected the change of crowding change by increase of cytosolic FRET ratio (Figure. 4C and 4D). The bright nature of CRONOS also enabled its spectrometric measurement in live cells, and we could successfully detect the increase of crowding by sorbitol (Figure. 4E and 4F).

As mentioned above, the FRET ratio of CRONOS passively diffusing into nucleolus was lower than that in nucleoplasm as well as in cytosol (Figure. 5A and 5B). On the other hand, the FRET ratio of CRONOS-NPM1 was higher than nucleoplasm (Figure.5C and 5D). NPM1 is reported to undergo LLPS with RNA and other proteins to form nucleolus in vivo ^26,27^. We thus hypothesized that the FRET ratio we obtained from CRONOS in the nucleolus reflected the molecular density of sparse nucleolar interstitium and not that of dense NPM1 droplets (Figure. 5E). If so, the CRONOS only exists in the interstitial space of nucleolus and it is not captured by the NPM1 droplets. To prove this, we adapted a simple model showing mobility of proteins spontaneously or passively locating to nucleolus using the fluorescence loss in photobleaching (FLIP) assay^28^. We overexpressed GFP-NPM1 and red fluorescent mCherry protein without targeting sequence in HeLa cells, followed by photobleaching of intranuclear mCherry signal. If a part of mCherry resides in the NPM1 droplets, remnant mCherry signal should be observed in the nucleolus area. After photobleaching, nuclear mCherry signal was quenched evenly, and no residual mCherry signal was observed in nucleolus, marked by GFP-NPM1 (Figure. 5F). After 10 sec of recovery, no significant accumulation of mCherry was observed in nucleolus either, showing that mCherry was not spontaneously uptaken by nucleolus. These data support the idea that the nucleolar CRONOS was not retained in the NPM1 droplets.

**Figure 5.**
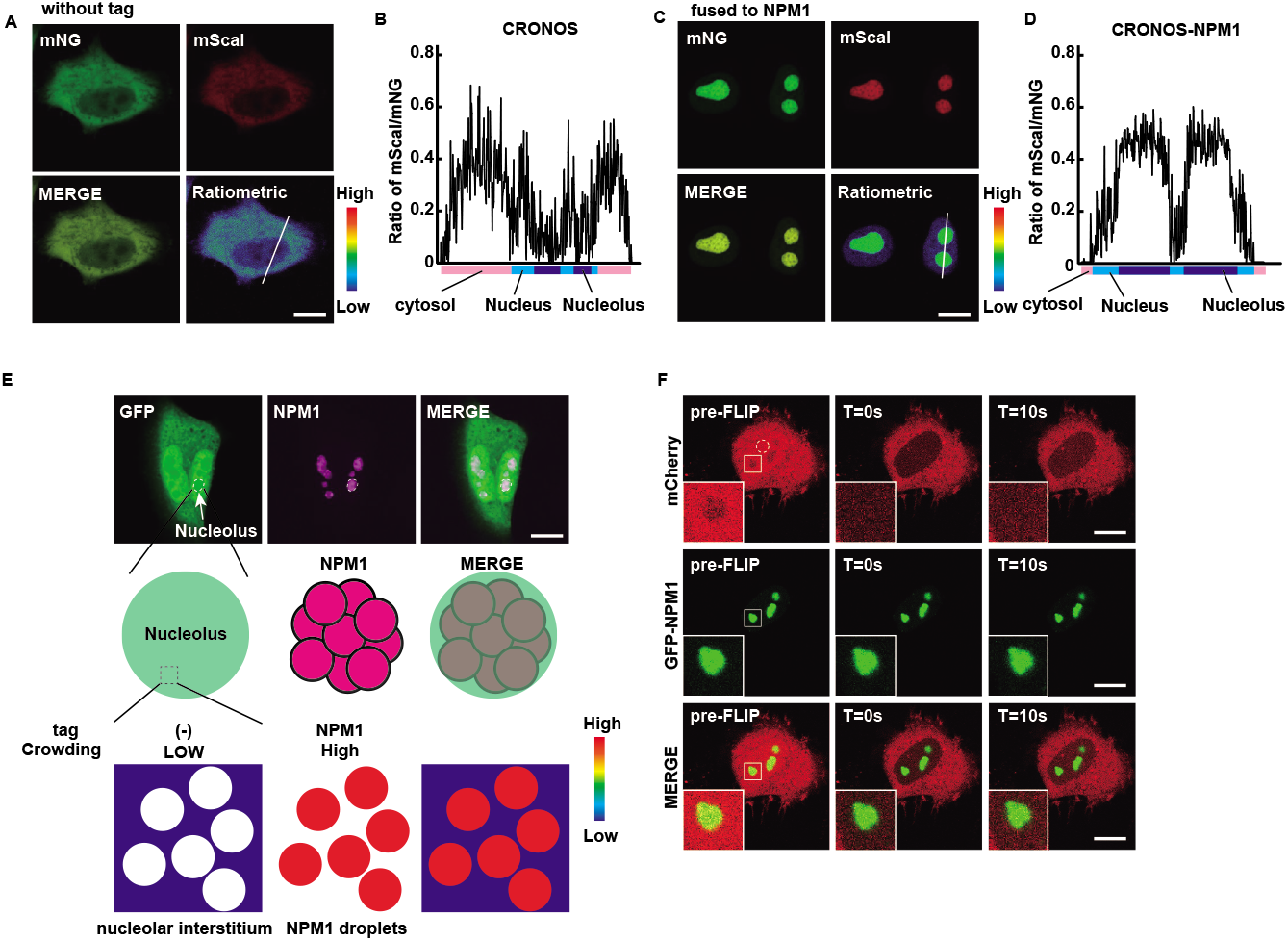
Uptake of CRONOS into nucleolus by tagging to NPM1 is necessary to assess the nucleolar crowding. (A) A confocal image of HeLa cells overexpressing CRONOS without tag. (B) The line profile of FRET ratio along the white line in (A). (C) A confocal image of HeLa cells overexpressing CRONOS-NPM1. (D) The line profile of FRET ratio along the white line in (C). (E) HeLa cells expressing GFP and DsRed-NPM1. When GFP is not tagged, GFP is supposed to localize to nucleolar interstitium. (F) FLIP analysis of HeLa cells expressing mCherry and GFP-NPM1. The circled area with a dashed line was bleached for long enough to quench nuclear mCherry fluorescence. The squared areas were zoomed. The scale bar shows 10 μm.

In light of these results, we further examined if CRONOS can sense the physiological change of crowding in MLOs. Nucleolus is one of the most important and largest MLOs in cells and the nucleolar architecture dynamically changes in response to various stress conditions ^29^. When the cells are exposed to extensive heat shock, the nucleolus reportedly gets condensed, confirmed by transmission electron microscopy^29^. On the other hand, treatment with actinomycin D, an anti-cancer compound which inhibits RNA polymerase activity, causes dispersion of NPM1 from nucleolus to nucleoplasm^6^. To examine the effects of environmental or chemical stress on nucleolus, the CRONOS-NPM1 sensor was expressed in HeLa cells, and the cells were exposed to heat shock (42°C for 3 hr) or actinomycin D (2 μg/ml for 3 hr) (Figure. 6A). The treatment with actinomycin D caused nucleolar disruption and the release of CRONOS-NPM1 from nucleolus to nucleoplasm with a little remnant in nucleolus. The FRET ratio from the sensor in nucleolus revealed that the degree of condensation was unexpectedly reduced by heat shock (Figure. 6B). The mCer-mCit also reported same tendency but with smaller changes (Figure S3A). The effect of heat shock on the localization of CRONOS-NPM1 was negligible, and the effect of temperature on the FRET ratio of recombinant CRONOS was also little (Figure S3B), suggesting that attenuation of rRNA synthesis by heat shock may explain the lower density of nucleolus^30^. The actinomycin D treatment strongly reduced the molecular crowding in nucleolus, also supporting the idea that repression of rRNA synthesis decrease nucleolar density. These results clearly indicate that CRONOS can detect changes in macromolecular crowding in the nucleolus.

**Figure 6.**
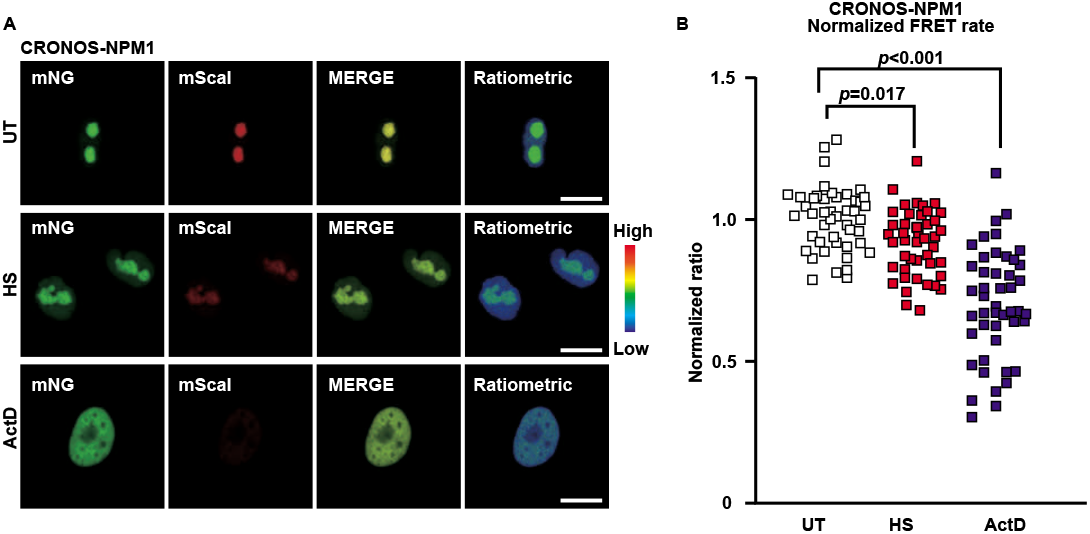
CRONOS-NPM1 successfully detects changes of molecular crowding in nucleolus under stress conditions. (A) Confocal images HeLa cells expressing CRONOS-NPM1 under untreated condition (UT) (upper panels), heat shock (HS) (middle panels) or treatment with Actinomycin D (ActD) (lower panels). Heat shock was done by incubation at 42°C. for 3 hr. Ratiometric images were pseudo-colored. (B) FRET ratio of CRONOS-NPM1. N=45 cells/condition. The scale bar shows 10 μm.

## Discussion

The emerging importance of LLPS and MLOs in biology provokes strong demand for technique to measure the biochemical features in the phase-separated droplets. In this study, we optimized the FRET pair of protein-based sensor, and successfully measured the macromolecular density of MLOs in live cells.

First, we sought for optimal FRET pair for creating a new crowding sensor with wider working range, enabling easier detection. The previous study reported the superior performance of mNG-mScaI pair for FRET^17^, and substitution of mCer to mNG as a FRET donor and mCit to mScaI as a FRET acceptor allowed us to create CRONOS, a novel crowding sensor with better performance. Furthermore, very bright nature of CRONOS also benefits for easier detection with better signal/noise ratio. Because mNG is one of the brightest green fluorescent protein^22^ and mScaI is one of the brightest red fluorescent protein^23^, CRONOS is much brighter than mCer-mCit sensor, enabling direct measurement of fluorescence in live cells with various modalities such as fluorescence spectrophotomer.

During the detailed characterization, we found out that both mCer-mCit sensor and CRONOS were not uptaken enough by the LLPS model droplets consisting of cationic peptides and ribonucleic acids, especially under existence of salt. The sensors were partially uptaken and distributed to the surface of the droplets, but not enough for accurate measurement, showing the difficulty to measure molecular crowding in the droplets by these sensors with their original shapes. In order to form LLPS, it is necessary for the constituent molecules to undergo multivalent interaction while complex correlations such as electrostatic interaction, cation-π interaction, and π-π interaction. The incorporation of the sensors were attenuated by the physiological concentration of salt, suggesting that phase-separated droplets consisting of the poly(PR) and RNA might uptake the sensor via electrostatic force. Because the measurement of single particle z-potential of the poly(PR) and RNA droplet revealed that the surface of the droplets are negatively charged^25^, this electrostatic force may play a central role in the accumulation of the sensor molecules to the surface of the droplets.

The advantage of protein-based sensor over nucleic acid-based one is its controllable delivery to specific region of interest by adding appropriate targeting signals^31^. When we fused the sensor to NPM1 or TIA1, we successfully delivered the sensor to different MLOs, nucleolus and stress granule. Although both nucleolus and stress granule are formed via LLPS, their ingredients are different and their crowding are also supposed to be different. Our study here revealed the previously undescribed difference of crowding among different MLOs: stress granules marked by TIA1 have much dense environment inside compared with nucleolus marked by NPM1. The discrepancy derived from the nucleolar FRET ratio obtained from CRONOS and that from CRONOS-NPM1 emerged the importance of uptake of the sensor into the phase separated droplets. As the FLIP analysis showed, the fluorescent protein without tag was not uptaken by MLO. Therefore, for measurement of crowding in MLOs, the sensor must be conjugated to the molecule which is a structural component of the MLO. Otherwise, the signals derived from interstitial space will lead to wrong interpretation. The crowding information from interstitial space is still useful because increase of crowding in the surrounding environment facilitates phase-separation of proteins. In other words, we need to measure crowding in both phase-separated droplets and interstitial space to fully understand the crowding status of MLOs.

Finally, we successfully detected changes of crowding in nucleolus under pathological conditions such as heat shock or inhibition of transcription by CRONOS-NPM1. Because nucleolus is the site for synthesis of ribosomal RNA, it was not surprising that suppression of transcription by heat shock or ActD treatment resulted in less-crowded nucleolus. The results were also confirmed with the mCer-mCit sensor, but CRONOS outperformed it in respect of dynamic range.

## Conclusion

In conclusion, we present here a novel crowding sensor CRONOS which allowed us to measure macromolecular crowding in MLOs in live cells. We found that CRONOS must be tagged to the molecule which undergoes LLPS to measure the crowding in the MLO of interest, otherwise CRONOS in the interstitial space will provide wrong information. CRONOS reported the previously undescribed differences of the degree of macromolecular crowding in different MLOs, and also detected changes of macromolecular crowding under physiological and pathological conditions. Further investigation of crowding in MLOs under various conditions and disease-related status will provide a novel insight into the pathophysiology of diseases caused by malfunction of MLOs.

## Acknowledgements

This work was supported by grants from the JSPS KAKENHI Grant numbers (16H06247, 17H03923, 20H03593, and 20H03827 to K. K., 19H03627 to N.S. and K.K., 20H02564 to Y. Hayamizu. and K.K., 17K15671 to Y. Harada, and 17H04067 to M. K.). This work was also supported in part by the Japan Agency for Medical Research and Development (AMED) (16ek0109180h0001 and 17ae0101016s0904), Strategic Research Foundation Grant-aided Project for Private Universities from the Ministry of Education, Culture, Sports, Science and Technology of Japan (M.K.), Takeda Science Foundation (K. K.), Japan Intractable Diseases (Nanbyo) Research Foundation (K.K.), the Tokyo Biochemistry Research Foundation (K.K.), and the Ichiro Kanehara Foundation (K.K.).

## Declaration of interests

The authors declare no competing interests.

## Supplementary Figures

**Supplementary Figure S1.**
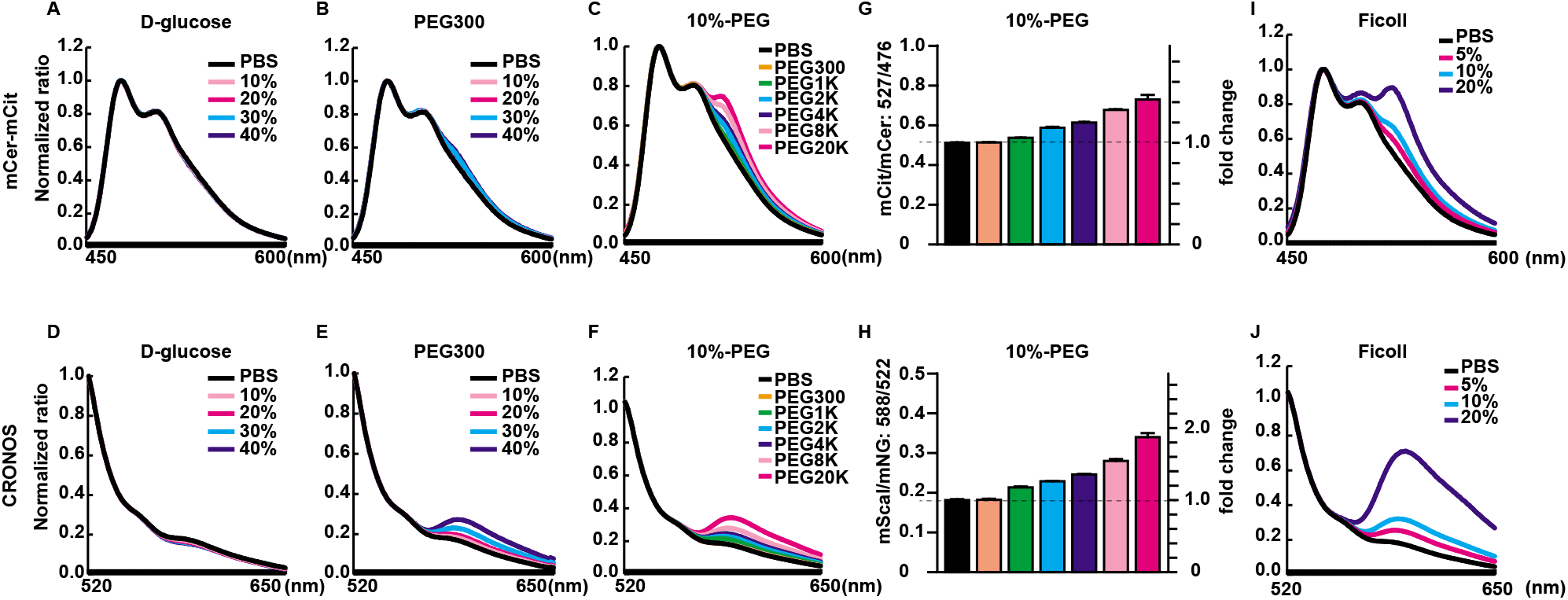
Fluorescence emission spectra of mCer-mCit and CRONOS reveal their response to various molecular crowders. (A)(B) Fluorescence emission spectra of the mCer-mCit sensor upon titration with 10%-, 20%-, 30%- or 40% of D-glucose or PEG300. The fluorescence emission spectra were normalized to the donor fluorescence. (C)(D) Fluorescence emission spectra of CRONOS upon titration with 10%-, 20%-, 30%- or 40% of D-glucose or PEG300. The fluorescence emission spectra were normalized to the donor fluorescence. (E) Fluorescence emission spectra of the mCer-mCit sensor exposed to 10% PEG with different molecular weight. (F) FRET ratio of the mCer-mCit sensor exposed to 10% PEG with different molecular weight. (G) Fluorescence emission spectra of CRONOS exposed to 10% PEG with different molecular weight were shown. (H) FRET ratio of CRONOS exposed to 10% PEG with different molecular weight were shown. The error bars indicate ±S.D. obtained from three independent measurements.

**Supplementary Figure S2.**
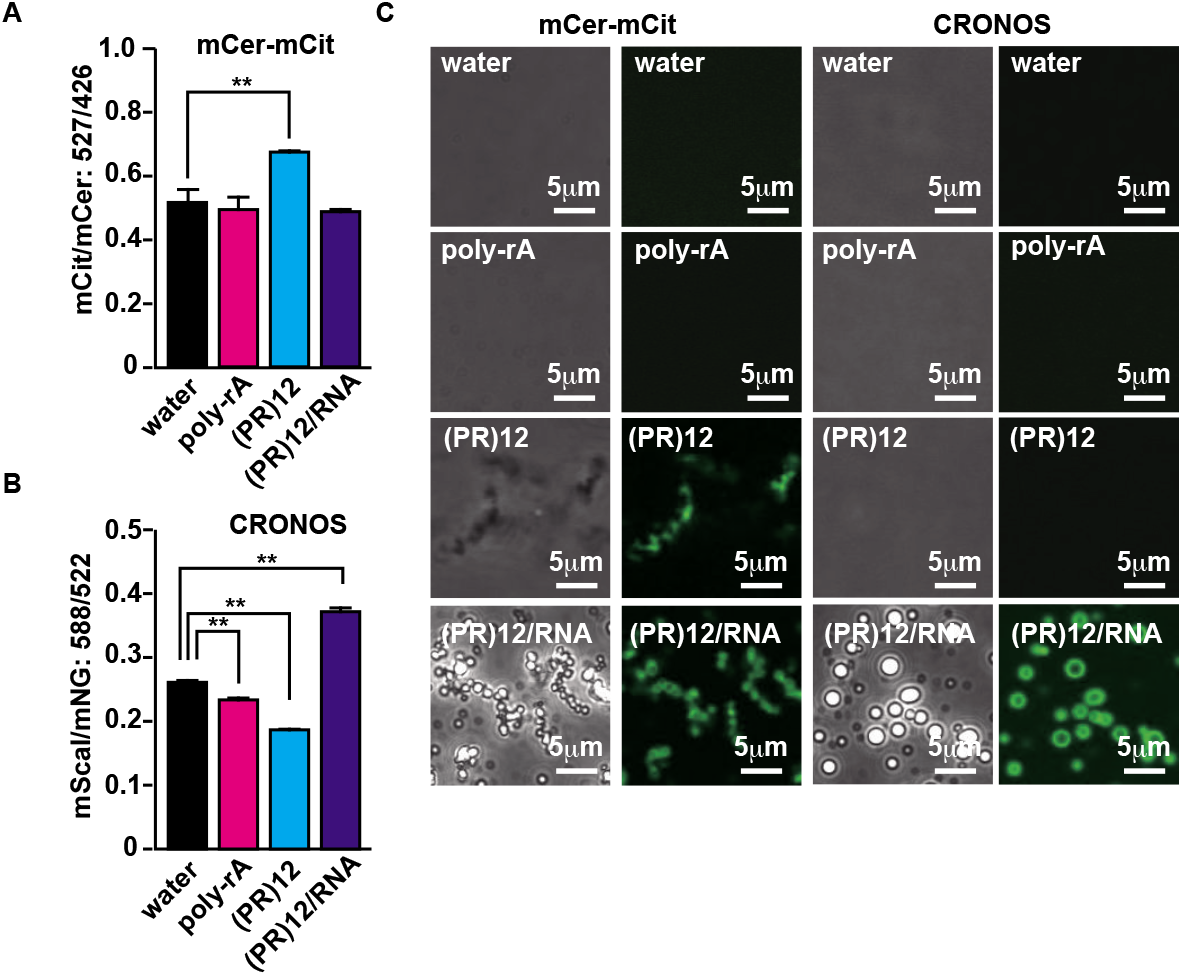
Uptake of crowding sensors into phase-separated droplets is insufficient in water. (A) The FRET ratio of the mCer-mCit sensor was not significantly changed whereas that of CRONOS detected molecular crowding in LLPS droplets in water. (B) Confocal imaging revealed that mCer-mCit sensor was weakly uptaken by the droplets and distributed to the surface of the droplets. CRONOS was also uptaken by the droplets and distributed to the surface.

**Supplementary Figure S3.**
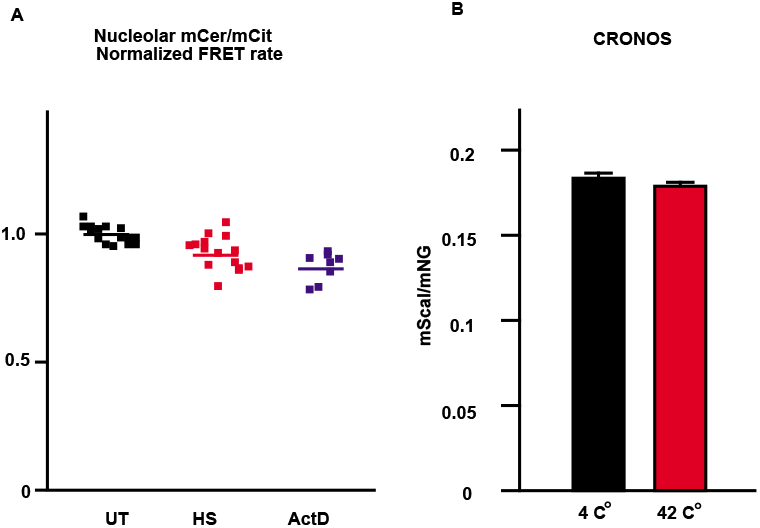
Heat shock and transcription inhibition changes nucleolar crowding. (A) The mCer-mCit sensor also detected changes of nucleolar crowding by HS or ActD treatment. (B) The temperature change does not affect the FRET ratio of recombinant CRONOS protein.

## Notes

### Competing Interest Statement

The authors have declared no competing interest.

